# MAGI: Mechanistic Consequences of Genetic Variants via Genomic Foundation Models

**DOI:** 10.64898/2026.05.31.729117

**Authors:** Dan Ofer, Stav Zok, Michal Linial

**Affiliations:** The Rachel and Selim Benin School of Computer Science and Engineering, The Hebrew University of Jerusalem, Jerusalem, Israel; Department of Biological Chemistry, The Life Science Institute, The Hebrew University of Jerusalem, Israel

## Abstract

Clinical variant interpretation requires mechanism-aware evidence to guide diagnosis and clarify the biological consequences of mutations. However, existing computational predictors and genomic foundation models largely function as black boxes, providing pathogenicity labels with limited mechanistic insight or clinical actionability. Here, we present MAGI (Mechanistic Annotation of Genomic Impacts), a novel method that bridges this interpretability gap by unifying clinically relevant variant interpretation with mechanistic genomic analysis. MAGI pipeline leverages a genomic transformer model to quantify the effects of DNA variants across 3,623 functional tracks, encompassing regulatory features, multi-omics datasets, including tissue specificity and chromatin states, and 21 additional molecular annotations of genes and transcripts. These signals are integrated through a deterministic logic layer that maps single-nucleotide variants and indels to explicit molecular consequences. We benchmark MAGI-derived consequences against clinical rationales curated from ClinVar and observe strong concordance that scales with the magnitude of functional disruption. MAGI accurately recapitulates canonical pathogenic mechanisms, including start codon loss, splice site disruption, and regulatory element perturbation, consistent with ClinVar annotations. We further present case studies addressing conflicting or incomplete mechanistic interpretations, as well as variants requiring complex inference. Notably, MAGI is also applicable to non-human genomes and was evaluated on multispecies OMIA pathogenic variants. Collectively, MAGI establishes a generalizable framework that extends beyond clinical diagnostics to enable mechanistic discovery in functional genomics, generating mechanistically grounded, testable hypotheses for variants of uncertain significance (VUS) and variants with discordant clinical interpretations. In several cases, MAGI proposes alternative explanations that challenge existing annotations, providing transparent rationales and experimentally tractable predictions.

## Main

“What does this genomic variation do?” remains a central question in genomics and clinical diagnostics. Most computational variant effect predictors frame interpretation as a binary classification problem. Although effective in aggregate, these approaches typically yield a single pathogenicity score with limited biological context. Over the past decade, numerous tools have been developed, spanning diverse methodological classes yet converging on a common inference paradigm. Widely used methods such as CADD ^1^, REVEL ^2^, SIFT ^3^, PolyPhen-2 ^4^. Much of the predictive signal arrives from evolution conservation ^5^, co-evolution based on sequence aligned positions ^6^, sequence context and functional annotations, often through supervised machine learning, to estimate deleteriousness in common ^7–9^ and rare variants ^10^.

Other specialized models, including SpliceAI ^11^ and AlphaMissense ^7^, focus on specific variant classes such as splice-altering and missense substitutions, respectively. Despite differences in architecture and scope, these methods generally compress complex molecular effects into a single internal score that is subsequently interpreted as a probabilistic or binary outcome (e.g., benign versus pathogenic) ^12, 13^. While highly effective for variant prioritization, this abstraction limits mechanistic interpretability ^14^.

There is therefore a growing need for approaches that translate predictive signals into explicit molecular consequences and perturbation studies ^14, 15^. Mechanistic rationales, such as disruption of regulatory elements, exon loss or transcript instability, are increasingly recognized as essential for clinical interpretation, assessment of medical actionability and experimental validation ^16^. Although experimental assays provide high mechanistic fidelity, they remain difficult to scale and cannot yet be comprehensively applied genome-wide ^12^. Recent advances in massively parallel reporter assays, deep mutational scanning and multiplexed functional genomics have highlighted both the power and the practical limitations of experimental variant characterization ^14, 17^, reinforcing the need for computational frameworks capable of mechanistic inference at genome scale ^8^.

Applied genome-wide, existing approaches largely fail to provide mechanistic explanations for variant effects beyond aggregate pathogenicity scores ^18^. To bridge this gap, we introduce MAGI (Mechanistic Annotation of Genomic Impacts), an unsupervised framework that extracts explicit molecular consequences from genetic variants using the Nucleotide Transformer v3 (NTv3) genomic foundation model ^19^. MAGI quantify the functional signal via the deviation (Δ) between reference and alternate alleles across 3,623 regulatory, functional, genomic, epigenomic, and other genomic-derived tracks ^20^. The extracted signals for the molecular alternation are then mapped via deterministic rules to produce human-interpretable molecular consequence annotations (e.g., “splice donor loss with intron retention” or “start codon disruption with ORF loss”). Rather than serving as a pathogenicity classifier, MAGI operate as a mechanistic annotator that is applicable across variant classes, including single-nucleotide variants (SNVs) and indels, and is uniformly deployed across the genome, including non-coding regions and expanding from human to other organisms.

## Results

### The MAGI framework

We illustrate the overall MAGI scheme in the context of human clinical interpretation. The MAGI framework is composed of two phases (**Fig. 1**). The first one is the implementation of the foundation model inference and the feature extraction (**Fig. 1a**). The difference between the model-predicted functional outputs for the reference (Ref) and alternative (Alt) alleles defines the functional deviation signal. The pipeline ingests Ref and Alt allele sequences centered in a 64 kb genomic window and processes them through the NTv3 650M model. Disruption scores (Δ) are computed across 21 structural genomic elements (BED format) and 3,602 epigenomic, tissue and cell line centric signals (BigWig format). Signals are mapped through magnitude and directional rules to generate rationales. The output of MAGI is a clinical interpretation for any observed genomic variants that explain the impact of such variants on the functionality of the affected unit.

**Figure 1.**
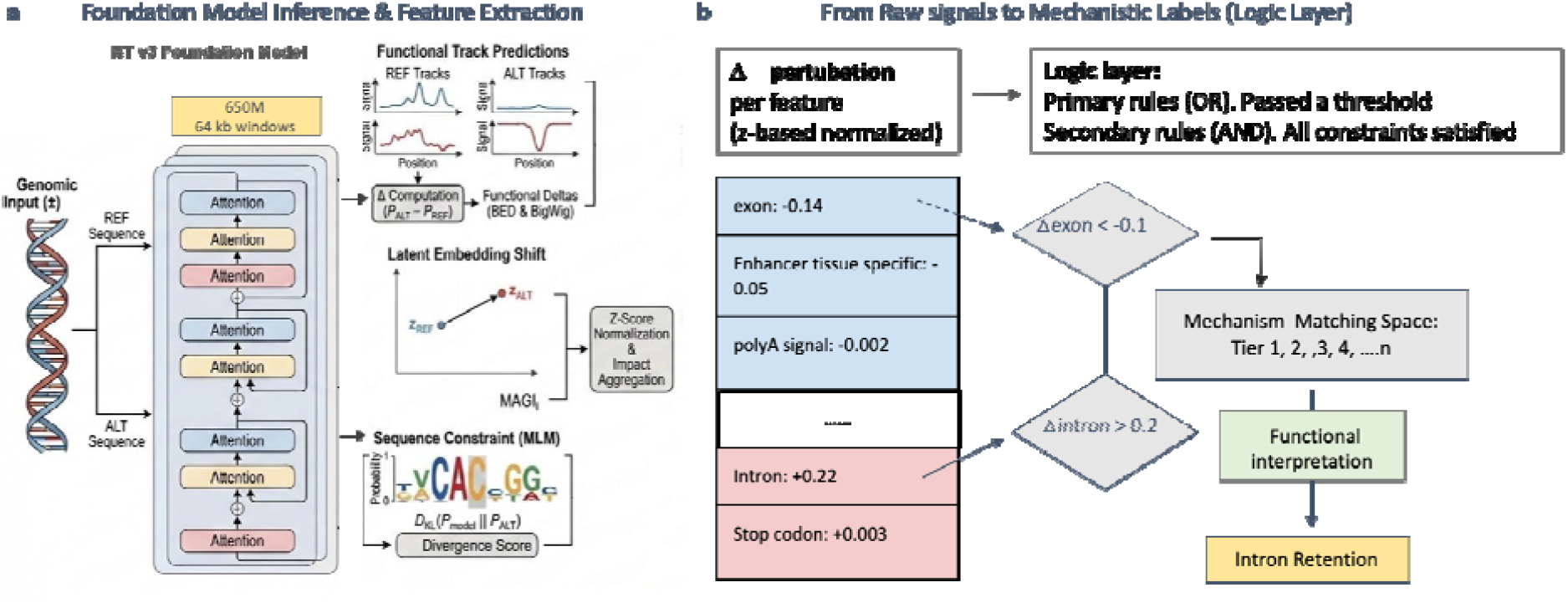
MAGI framework for mechanistic annotation of genomic variants. **(a)** Input representation and effect quantification. Single-nucleotide variants (SNVs) and indels are encoded by constructing paired reference (Ref) and alternate (Alt) sequences centered on the variant locus, preserving local genomic context. Both sequences are processed by the Nucleotide Transformer v3 (NTv3) to generate predictions across up to 3,623 functional tracks. Variant impact is quantified as the difference (Δ) between Alt and Ref signals, capturing perturbations in regulatory, epigenomic, and sequence-level features (hereafter, MAGI score). **(b)** Mechanistic inference via logic layer. Δ signals are integrated with gene and transcript annotations and resolved through a deterministic logic layer that assigns explicit molecular consequences, including coding, splicing, and regulatory effects, yielding interpretable, mechanism-based variant annotations.

An additional branch of MAGI incorporates language-model-derived metrics, including log-likelihood ratios (LLRs), to quantify how variant-induced sequence changes alter predicted probability distributions. In this framework, differences between reference and alternative sequence contexts can be evaluated using measures such as Kullback–Leibler (KL) divergence. MAGI applies sequence-constrained masked language modeling, in which predictions are generated under explicit constraints imposed by the underlying biological sequence. This enables the detection of variant-associated shifts in regulatory or binding-related sequence patterns, including evidence involving transcription factors (TFs) and DNA/RNA-binding proteins (**Fig. 1a**).

For MAGI’s second phase, we extracted rule-based logic from the available molecular and genomic data to capture relationships between DNA sequence variation and transcripts, open reading frames (ORFs), splicing, cis-regulatory elements, and MLM-derived constraints. This rule-based framework is designed to generate both primary and secondary decision rules, allowing MAGI to move beyond single-feature interpretation toward richer, combinatorial logic using AND/OR relationships (**Fig. 1b**).

Candidate consequences are then prioritized and assigned to a tiered decision structure. These tiers include, for example, definitive loss-of-function (LoF) events, splicing alterations, exon-structure disruptions, regulatory-element perturbations, and additional mechanistic categories. Together, the tiers capture complementary dimensions of variant impact and provide a structured framework for ranking biologically meaningful candidates (**Fig. 1b**).

MAGI predictions are then evaluated against clinical annotations from resources such as ClinVar ^21^ and OMIA ^22^ across human and non-human species, respectively. Within the MAGI-ClinVar validation scheme, we use LLM to judge on the levels of agreement between the detailed rational reported in ClinVar and the outcome of MAGI (**Fig. 2a**). MAGI agreement level is not based on the label of pathogenicity for a variant but on Phase 1 NT-v3 signals (genomic annotation, epigenomic/transcriptional, and sequence-context dimensions), and current knowledge. Recall that ClinVar rationale provides beyond the variant pathogenic classification (SNP or indel), the entire information known from SNP databases, experimental validation and publications that support the inference. Along with the evaluation task, the degree of concordance between predicted mechanisms and known pathogenic rationale is recorded with full match (concordance), partial match, and discordance (**Fig. 2a**).

**Fig 2.**
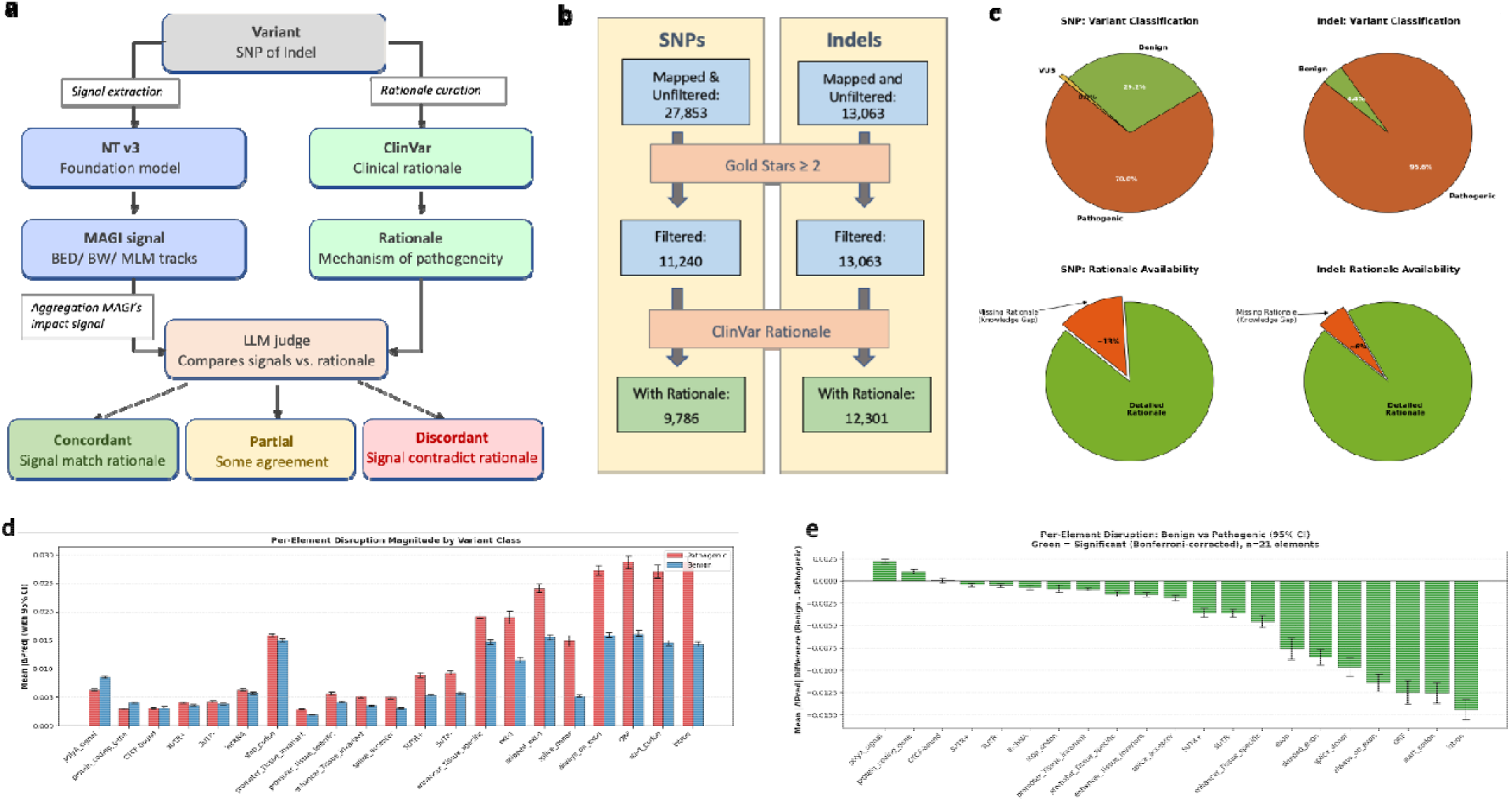
MAGI variant processing and performance over ClinVar variants. **(a)** Evaluation of NTv3 derived signals against the ClinVar rationale using LLM as judge. The agreement levels are partitioned to concordant, partial and discordant. **(b)** Total number of detected variants following genome alignment (hg38), separated into single-nucleotide polymorphisms (SNPs) and insertions/deletions (indels). The initial number of SNPs is ~40k SNPs that could not be uniquely mapped to the hg38 version. Variants with supporting clinical interpretations are labeled ClinVar rationale, with confidence stratified by review status (≥2 gold stars). The gold stars ≥2 filter removed SNPs that have not been supported by at least two independent submitters. Quality control and filtering steps applied to remove low-confidence and technical artifacts, yielding a high-confidence variant set (marked as filtered). Note that this step did not filter any of the mapped indels. SNPs that were maintained after removal of those lacking ClinVar rationale are used to evaluate the accuracy and precision of the MAGI approach. **(c)** Variant classification distribution by ClinVar for SNPs (left) and Indels (right). ClinVar identifies 70% of the SNPs with rationale (n=9,786) to be pathogenic and the rest are benign. For a small fraction (0.9%), variants labelled as uncertain significance (VUS). Among the indels, 95.6% are assigned as pathogenic with a minimal number of indels labelled as benign. ClinVar rationale covers only 87% of the confident SNPs, leaving a substantial gap in knowledge. For indels, only 6% do not have rationale among the supported ClinVar indels. **(d)** Per-element disruption by variant class. Distribution of the 21 BED element delta scores (Δ) for pathogenic versus benign variants, stratified by molecular consequence class in Clinvar. **(d)** Per-element disruption by variant class. Distribution of the 21 BED element delta scores (Δ) for pathogenic versus benign variants, stratified by molecular consequence class in Clinvar. Statistical significance for the median by Mann-Whitney U p-value, and rank by the effect size r. **(e)** Mean delta prediction of benign vs pathogenic label according to the 21 BED elements in ClinVar with 95% of CI. Colored green all significant elements (Bonferroni corrected). The order of the elements is identical for d and e.

### Variant curation from ClinVar

To test MAGI performance, we extracted a gold-standard dataset of expert-curated variants composed of both indels and SNPs, annotated by ClinVar as Benign, Pathogenic, or VUS, most of which also included free-text clinical justifications from ClinVar (**Fig. 2b**). The SNP set included 9,786 variants with rationale out of 11,240 filtered SNVs, while the indel set included 12,301 variants with rationale out of 13,063 filtered indels. Variants were retained only if they had a ClinVar review status of two stars or higher, ensuring a relatively high-confidence benchmark set.

Within the SNV cohort, the majority of variants were classified as Pathogenic (70.8%), followed by Benign (29.2%), with only a very small fraction labeled as VUS (**Fig. 2c**). In the indel cohort, the distribution was even more skewed toward pathogenicity, with 95.6% Pathogenic and 4.4% Benign variants. Notably, rationale availability was high in both groups, covering 87.1% of SNVs and 94.2% of indels, thereby enabling a large-scale comparison between MAGI-derived molecular signals and expert clinical interpretation (**Fig. 2c**). This curated benchmark provided two complementary evaluation layers: (i) The ability of MAGI-derived features to distinguish known benign from pathogenic variants. (ii) The extent to which the molecular signals inferred by the model were concordant with the mechanistic interpretation described in ClinVar free-text rationales.

### Functional element disruption scales with variant consequence

To assess whether MAGI’s predictions capture bona fide molecular mechanisms, we stratified variants by annotated consequence and examined the magnitude of disruption in the corresponding genomic features. Across all categories, pathogenic variants exhibited significantly greater perturbations than benign variants in the expected elements (**Fig. 2d**). Start codon loss showed the most pronounced separation, with a median Δ_start_codon of 0.36 for pathogenic variants compared to 0 for benign variants. ORF disruption yielded the largest overall effect size (rank-biserial r = 0.632, p < 1.0×10^−99^, n > 10,000), followed by exon loss (r = 0.557, p < 1.0×10^−78^). Splice-related features (derived from SpliceAI) were consistently significant, including splice acceptor (r = 0.120, p = 0.025), intronic regions (r = 0.443, p < 1.0×10^−16^), and exonic regions (r = 0.495, p < 1.0×10^−20^). Variants annotated as missense exhibited more modest yet detectable disruptions in regulatory tracks (r = 0.328, p < 1.0×10^−40^), consistent with their primary effects occurring at the protein level rather than through genomic regulatory elements. **Fig 2e** displays the statistically significant (Bonferroni corrected) mean difference between benign and pathogenic signals across molecular elements. The analysis considers the 21 molecular annotations.

Together, these results support a coherent hierarchy of MAGI-derived signals: structural disruptions (e.g., start codon loss, ORF disruption, exon loss) produce the strongest effects, splice-related perturbations are intermediate, and missense variants are not explicitly detectable, reflecting their distinct mechanistic basis.

### Population-aware exoneration exposing variants with incomplete penetrance

Mapping the MAGI Impact Score against gnomAD allele frequency (AF) shows a footprint of purifying selection. Variants with high impact scores are rare, consistent with negative selection against functionally disruptive variants. The information captured in the MAGI Impact Score allowed it to reach an ROC-AUC = 0.69 for pathogenic vs. benign separation and overall effect size (measured by Cohen’s d) of 0.65 (**Fig. 3a**). This dual-axis approach enables the systematic exoneration of variants. Alleles that exhibit high mechanistic impact but show high frequency in specific subpopulations can be flagged as tolerated, buffering against false positives and highlighting sites of potential compensatory regulation, or traits affecting large population subgroups.

**Fig 3.**
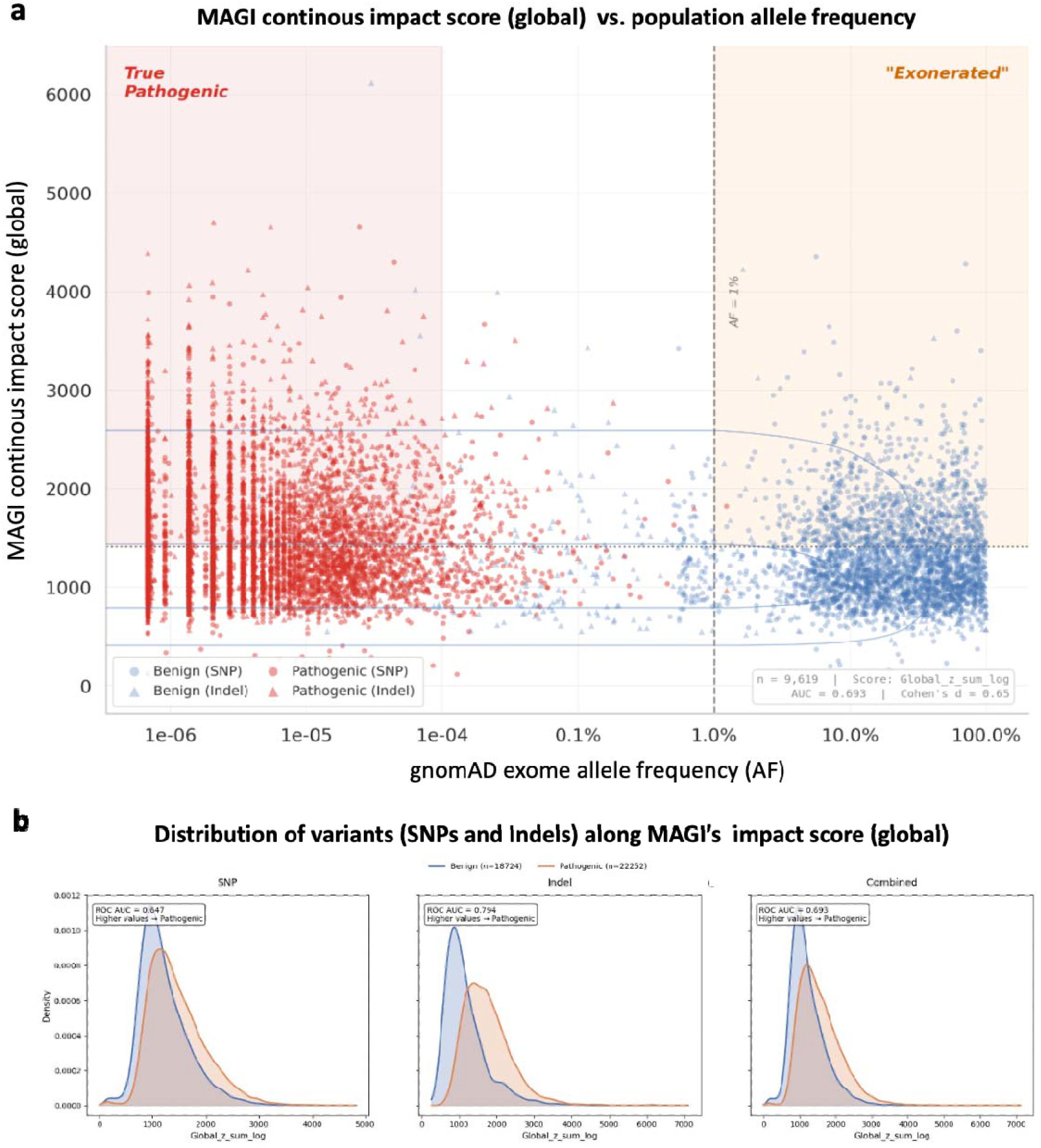
MAGI score in view of AF in the population. **(a)** Scatter plot of the MAGI Impact Score versus gnomAD allele frequencies (AF). For each variant, the MAGI score is a real-valued burden score where higher values indicate total functional deviation between Alt and Ref predictions. This global sum of the z-score delta tracks in log (see Methods) is plotted and not a probability. No fixed pathogenic cutoff is assumed. The horizontal line in the plot is the median pathogenic score used as a visual reference only. The allele frequency >1% and <0.01% are marked for visualization. **(b)** Distribution of MAGI continuous impact scores for SNPs and indels classified as benign or pathogenic. Density plots show the distribution of the aggregated MAGI impact score (Global_z_sum_log). Higher MAGI scores correspond to larger predicted functional perturbations derived from cumulative deviations between Ref and Alt alleles. Pathogenic indels (middle) display a marked rightward shift toward higher impact scores relative to benign indels, indicating stronger predicted molecular disruption (ROC-AUC 0.794). Density estimates were generated using kernel smoothing, with shaded regions representing the probability density for each class.

This analysis also indicates that a critical confounder in variant interpretation is incomplete penetrance, where variants driving measurable molecular disruptions fail to manifest as disease. Unimodal computational predictors frequently over-call these variants as pathogenic. To contextualize MAGI’s predictions within human populations, we integrated the log-transformed sum of normalized functional deviations (MAGI Impact Score) with gnomAD population allele frequencies. We confirm a large depletion of high-impact variants at higher frequencies (purifying selection) and isolate the exonerated variants, those with profound predicted molecular changes that are tolerated in the population.

Using the entries set of 40k unfiltered (but mapped to hg38) variants composed of 46% labelled as benign and 54% as pathogenic showed a stronger separability for the Indels analysis (**Fig. 3b**). Restricting the analysis to the SNP set, yielded an ROC-AUC of 0.647 but the signals related to the indels reached ROC-AUC of 0.794. We will not discuss the few variants that show a surprising trend such as AF >1% with pathogenic label and a few instances of benign variants with AF <1.0e-04, and a high MAGI Impact Score. The separation between the two distributions demonstrates that MAGI effectively capture pathogenic signals for insertion/deletion variants despite their heterogeneous size and sequence context.

### MAGI concordance with ClinVar rationale improves with model-derived confidence

We next compared MAGI-derived mechanistic annotations against a gold-standard benchmark of expert-curated free-text clinical rationales from ClinVar to assess whether model-derived molecular signals were mechanistically concordant with human interpretation. Across the curated benchmark, MAGI showed substantially stronger performance for SNPs than for indels (**Fig. 4a**). Among SNPs, 78% of variants were fully concordant with ClinVar rationale and an additional 9% were partially concordant, yielding 87% overall agreement. In contrast, indels showed markedly lower performance, with 18% fully concordant and 25% partially concordant interpretations (43% overall agreement), highlighting the greater difficulty of capturing indel consequences using sequence-derived functional signals alone.

**Fig 4.**
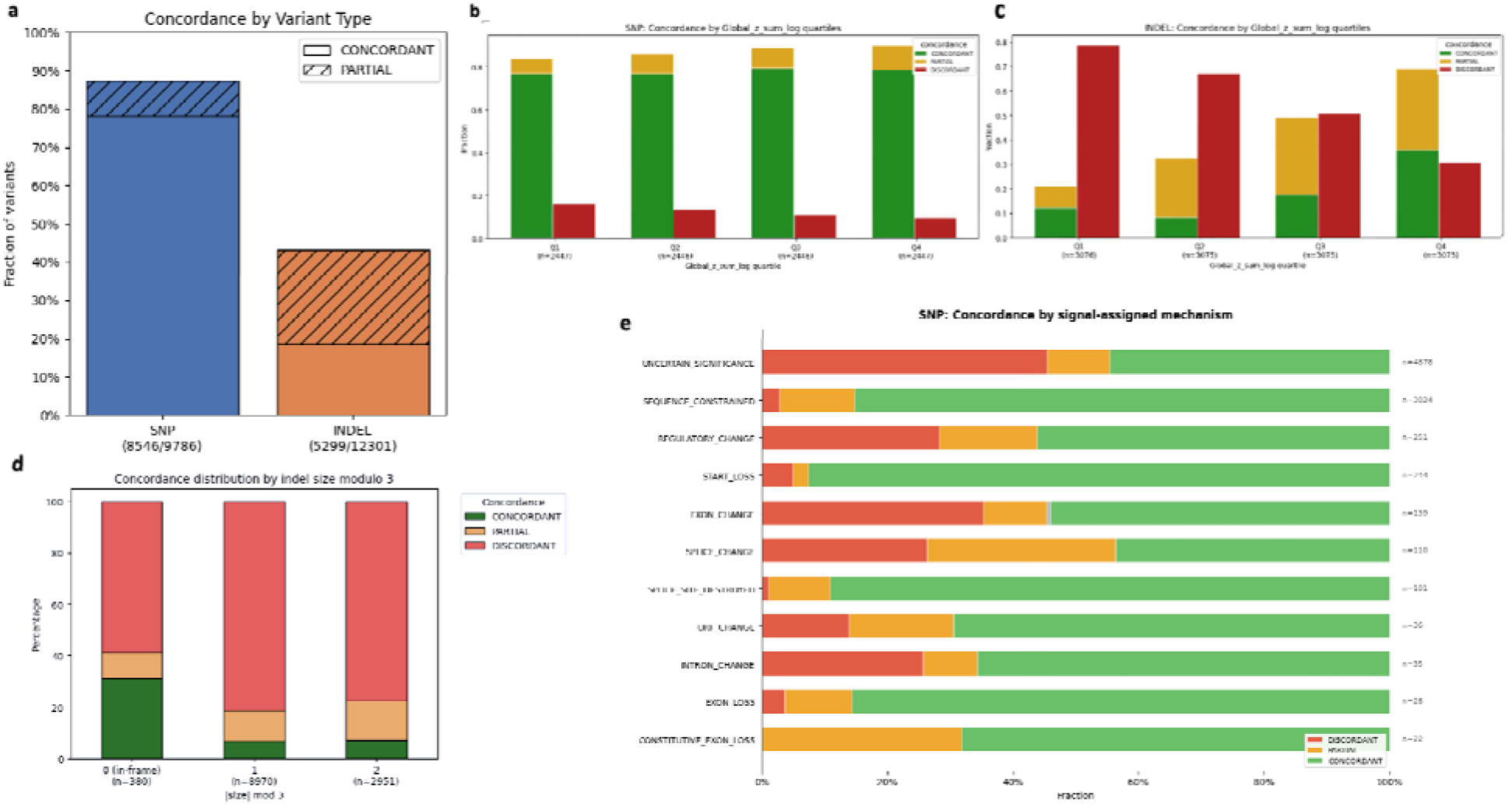
Concordance by variant type, MAGI-score and assigned mechanisms. **(a)** Concordance of MAGI-derived mechanistic annotations with curated ClinVar rationale. Variants for which MAGI fully agrees with the supporting clinical interpretation are labeled concordant, whereas partial indicates variants for which MAGI captures part, but not all, of the ClinVar rationale. Overall concordance (concordant + partial) reaches 87% for SNPs and 43% for indels. **(b)** Concordance across quartiles of the MAGI score for SNPs. Concordance remains high across all quartiles, with a modest improvement toward higher-scoring variants. **(c)** Concordance across quartiles of the MAGI score for indels. Discordance decreases markedly with increasing MAGI score, accompanied by a monotonic rise in overall concordance. **(d)** Indels’ concordance according to the frameshift caused by the insertion/deletion. Only the indels divided by 3 (marked as in-frame) reach 40% concordance while the other shifts agree with ClinVar at <20%. **(e)** Concordance of MAGI-derived mechanistic annotations across SNPs with assigned mechanisms. ‘Uncertain significance’ denotes variants whose model-derived signals were too weak to exceed MAGI’s rule-based annotation thresholds. ‘Sequence constrained’ denotes variants for which the strongest signal was derived from language-model features (e.g., Alt vs. Ref likelihood ratio). High concordance is observed for categories associated with well-defined molecular disruption, including start-loss and splice-site alterations.

To determine whether these predictions reflected meaningful biological perturbation rather than low-level background signal, we stratified concordance by quartiles of the MAGI score, our aggregate measure of model-derived functional disruption. For SNPs, concordance was already high across all quartiles, with a modest but consistent increase toward higher-scoring variants (**Fig. 4b**). In contrast, indels showed a strong monotonic trend, with discordance decreasing substantially and overall concordance rising from 21% in the lowest quartile to 69% in the highest quartile (**Fig. 4c**). Together, these results indicate that stronger MAGI-derived signals are associated with greater mechanistic agreement with curated clinical interpretation, particularly for variant classes that are intrinsically harder to resolve.

For indels, concordance also depended somewhat on whether the event was in-frame or frameshifting (**Fig. 4d**), consistent with the idea that different indel classes generate distinct molecular consequences that are not equally captured by the current framework. Finally, among SNPs for which MAGI assigned a specific mechanism, several categories showed particularly high agreement with ClinVar rationale, including sequence-constrained, start-loss, and splice-site alterations (**Fig. 4e**), indicating that MAGI is most reliable when the predicted signal aligns with well-defined molecular consequences.

### Assisting clinical utility by MAGI inference

A primary objective of computational variant annotation is to aid clinical decision making. We applied MAGI to clinical variants characterized by missing annotations or discordant classifications in external databases to test its utility as an engine for hypothesis generation. In clinical databases, variants may be labeled “pathogenic” without documented mechanisms. Many of these are labeled on the basis of their rarity. For example, a rare MAT1A classified as a VUS by ClinVar lacked a clinical rationale (**Fig. 5a**). MAGI autonomously identified, in addition to the missense mutation, a transcript-level perturbation. Furthermore, in cases where MAGI predictions strongly conflict with “benign” labels, the model frequently identifies severe, localized regulatory disruptions (e.g., ectopic enhancer creation). These discordant flags provide precise molecular hypotheses, guiding targeted functional validation for variants that may exhibit tissue-specific or incomplete phenotypes missed by standard diagnostics. We will present several case studies, each highlighting a distinct dimension of MAGI inference.

**Figure 5.**
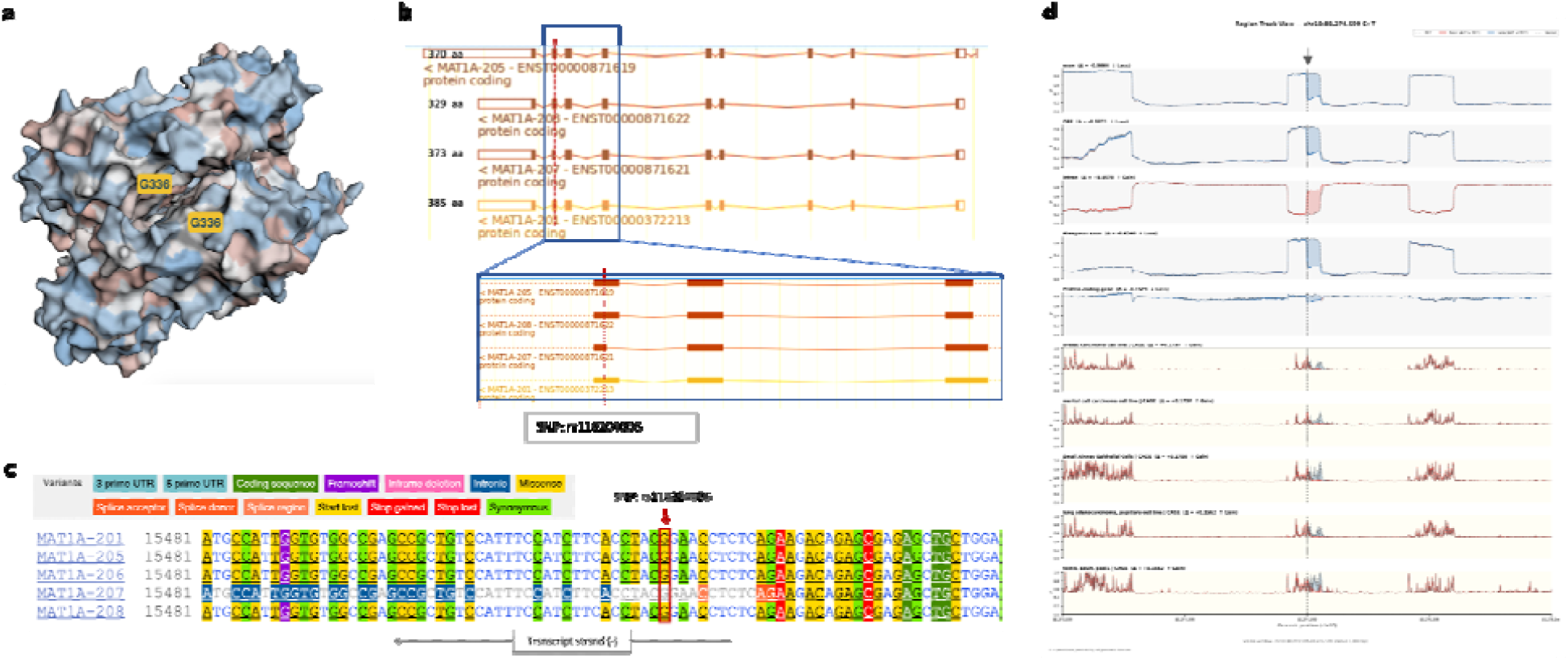
Multi-level impact of the MAT1A p.Gly336Arg variant across protein structure and transcript isoforms. **(a)** Structural model of MAT1A highlighting the position of Gly336 within the C-terminal domain. The results of the Missense3D approach indicated a disruption of the buried glycine (position 336). The model is based on PDB: 6SW6 dimer of a full length MAT1A. **(b)** Transcript models showing the genomic position of the variant across MAT1A isoforms, with the red dashed line indicating the variant position in full transcripts and a zoom in to exons 7 to 9 in the representative variants. **(c)** Sequence alignment across transcripts, with the variant position marked (red frame, arrow), illustrating nucleotide context interpretation to known variants. Note that MAT1A-207 may influence splicing regulatory elements with splicing donors used differently than the canonical one (e.g., in variant MAT1A-201). **(d)** MAGI transcript-level tracks for rs118204006. Genome-wide tracks show predicted functional changes around the variant. Blue regions indicate predicted loss, red regions indicate predicted gain of signal. Tracks include exon and intron (~400 bp), Δ scores, ORF and coding gene Δ, and tissue-specific transcription activity derived from CAGE. The vertical line marks the position of rs118204006. Recall that the transcript is on the reverse strand.

### MAGI predicts variant p.Gly336Arg in MAT1A to impact transcript processing

The MAT1A gene encodes methionine adenosyltransferase, a key hepatic enzyme responsible for the synthesis of S-adenosylmethionine (SAM), the principal methyl donor in cellular methylation reactions. Through this central role, MAT1A is essential for methylation-dependent metabolic pathways, and its disruption leads to methionine adenosyltransferase deficiency (MATD) and impaired methionine metabolism. The variant rs118204006 (NM_000429.3:c.1006G>A; p.Gly336Arg), located at chr10:80274599 (GRCh38), results in a non-conservative amino acid substitution within the C-terminal domain of the protein. It is extremely rare in the general population (reported on 11 alleles among >800,000 individuals in gnomAD), supporting a potential disease-associated role. Multiple in silico tools predict a deleterious effect, and functional studies demonstrate reduced enzymatic activity to ~23% of the reference wild-type, consistent with a partial LoF mechanism. Clinically, the variant has been reported in compound heterozygous states, producing conflicting ClinVar classifications as variant of uncertain significance (VUS) with another report labels the same variant as likely pathogenic ^23^.

Structurally, Gly336 is located in a buried region near the dimerization interface (**Fig. 5a**), suggesting that substitution to a bulky, charged arginine residue may destabilize local packing or interfere with dimer formation. This is consistent with experimental observations of reduced enzymatic activity ^23^. At the transcript level (**Fig. 5b**), MAT1A exhibits multiple isoforms (14 transcripts among them 12 protein-coding isoforms), and the impact of the variant appears isoform-dependent. Notably, in transcript MAT1A-207, the variant lies in proximity to an alternative exon–intron boundary, where it may influence local splicing regulatory elements. Sequence alignment across transcripts (**Fig. 5c**) highlights context-dependent differences, suggesting that the variant could perturb exonic splicing or nearby sequence motifs (e.g., pyrimidine-rich regions), potentially altering exon inclusion or splice efficiency. MAGI revealed effects extending beyond protein-level disruption (**Fig. 5d**). Genomic tracks surrounding rs118204006 showed a pronounced loss of exon signal (blue) accompanied by a gain of intronic signal (red), consistent with altered transcript processing. Integration of large-scale experimental tracks (BED-derived scores) further indicated coordinated regulatory perturbations, including reduced local expression across multiple tissues and cell lines. Together, these findings support an overlooked transcript-level mechanism in addition to the established enzymatic defect.

Altogether, the data suggest a dual pathogenic mechanism for p.Gly336Arg, combining partial loss of enzymatic activity with altered transcript processing. This case highlights the limitations of binary pathogenicity classifications and underscores the value of integrative, isoform-aware mechanistic interpretation. Although ClinVar classifies this variant as either VUS or likely pathogenic, the combined structural and transcriptomic evidence points to a more nuanced mechanism involving both protein destabilization and splicing disruption. Given the variable phenotypic consequences of MAT1A mutations, including recessive and dominant-negative effects, and the further reduction in enzymatic activity observed in compound heterozygous states (e.g., with p.Arg264Cys), this example illustrates how MAGI can uncover multi-layered molecular consequences by integrating allelic and transcriptomic context.

### MAGI predicts the role of a SNP in ALOX15B in disruption of tissue-specificity through alternative splicing

We analyzed rs9895916, a common SNP (global MAF ~20%) associated with a missense variant in ALOX15B (Arg486His; NP_001132.2, transcript NM_001141.3). This variant is not reported in ClinVar, and classified as benign by VarSome. ALOX15B encodes a lipid-oxidizing enzyme distinct from ALOX15, preferentially metabolizing arachidonic acid and exhibiting singular specificity for DHA (docosahexaenoic acid). Its expression increases during monocyte-to-macrophage differentiation and is further modulated by cytokines and hypoxia. Functionally, ALOX15B function is often tissue and cell specific. For example, it contributes to hypoxia responses in the heart, affecting cardiomyocyte activity, and regulates cholesterol metabolism, inflammatory signaling, and T-cell migration in macrophages.

**Fig. 6a** shows the MAGI confidence for the unaltered and altered nucleotide (marked Ref and Alt). It highlights the strong signal derived from BED tracks. The rs9895916 variant is predicted to markedly alter splicing signals, with increased intronic assignment and reduced ORF and exon definition. **Fig. 6b** presents sequence alignment of seven annotated transcripts. In four transcripts, the variant lies at a splice donor site, whereas in the remaining three it resides within an intronic region. **Fig. 6c** illustrates the genomic view (focus on 2000 nt) from MAGI application. The tracks combine BED (top) and BigWig (bottom) tracks. The variant influences expression likelihood via altered splicing outcomes. **Fig. 6d** shows tissue-specific alternative splicing patterns (GTEx) for transcripts capturing the relevant junctions, with junctions 11, 12, and 13 predicted to be affected by the variant change. We conclude that MAGI can predict even a relatively minor impact of rebalancing the expression level of the different variants and changes in the tissue specificity by altering alternative splicing outcomes.

**Figure 6.**
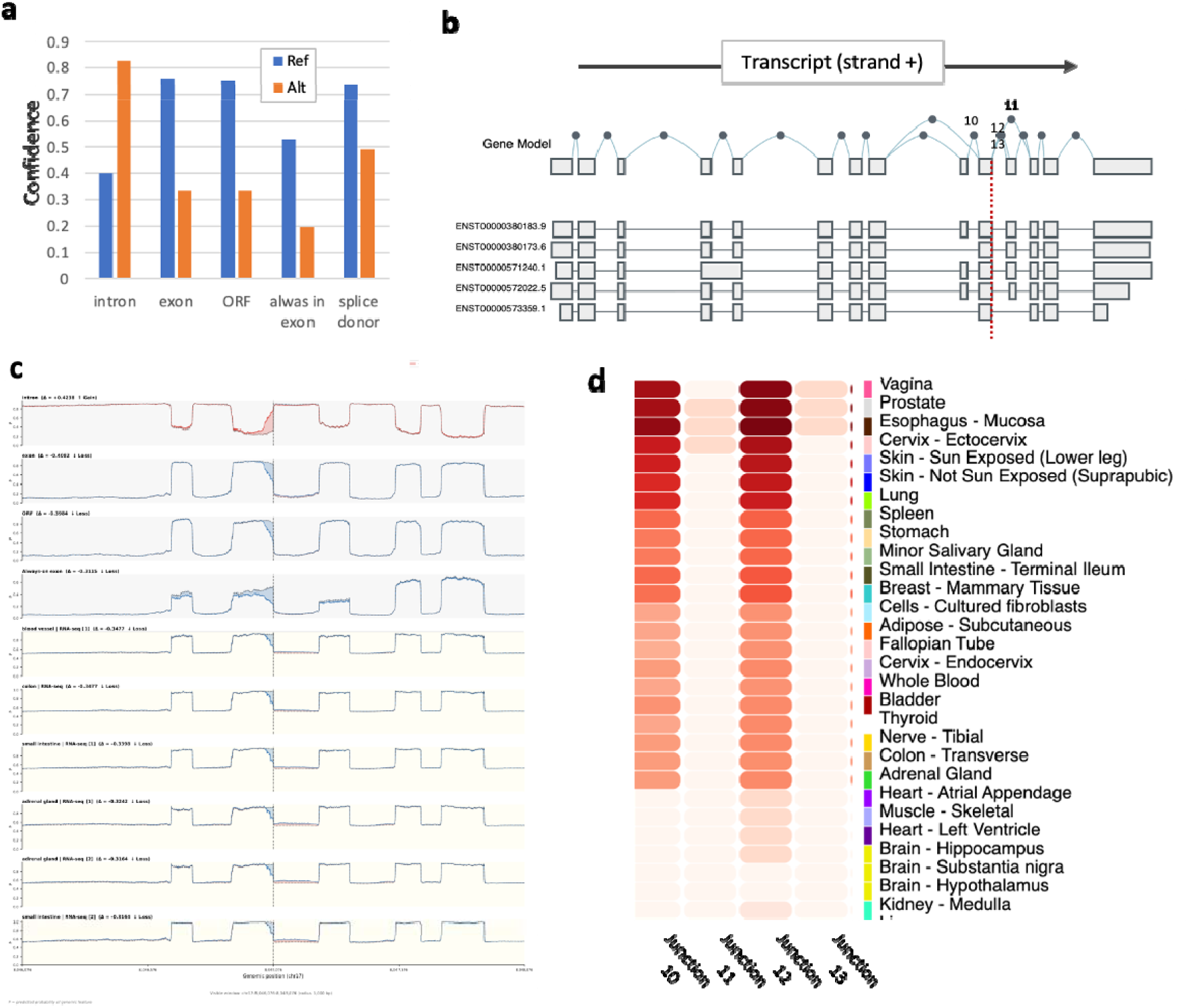
Overview of ALOX15B transcript structure, sequence variation, and expression. **(a)** Bar plot showing confidence levels calculated by MAGI across BED track annotations for the reference (Ref) vs. alternative (Alt) alleles. **(b)** Schematic of ALOX15B gene models and transcript isoforms. The vertical dashed line marks the position of the SNP. Junctions 11-13 involved the apparently impaired donor site by the SNP. **(c)** MAGI prediction track-centric view highlights the changes in exon likelihood at the vicinity of the variant. **(d)** Heatmap displaying relative counts identified per splicing event 10-13 (according to the schematic view in b) across tissues from GTEx. Junctions 11 and 13 are used only in selected tissues (e.g., prostate), whereas junction 12 shows broader usage across tissues.

The GLA c.801G>A (p.Met267Ile; ChrX: 101398785 C>T) variant has been documented in ClinVar with consistent evidence supporting its pathogenicity in Fabry disease (OMIM accession no. 301500). Among six submissions, two classify it as pathogenic and four as likely pathogenic. Notably, in a hemizygous male patient, this variant correlates with markedly reduced enzymatic activity of α-galactosidase A, the biochemical hallmark of Fabry disease. Further support indicated other missense substitutions in Met267 that were reported in affected individuals. However, GLA c.801G>A variant is absent in large population datasets, including 807,000 individuals in gnomAD v4.1.1, reinforcing its rarity and clinical relevance.

Using MAGI, we obtained complementary mechanistic insights that expand on this established understanding. Unlike traditional interpretations focused solely on amino acid (aa) substitution, MAGI predict a strong splicing impact for Met267Ile. The probability-delta score indicates significant disruption of a canonical splice donor site at the exon-intron boundary (Δ splice donor = −0.529 versus reference allele = 0.736; LLR = −1.056). This suggests that the variant may lead to aberrant splicing, potentially producing truncated or misprocessed transcripts. Importantly, this predicted splicing defect provides a mechanistic explanation that is independent of direct effects on the α-galactosidase biochemical activity.

**Fig. 7a** analyzed all 17 transcripts reported for GLA gene (ENSG00000102393). While several transcripts are likely to produce proteins of different length, others are expected to be subjected to NMD degradation. Still in 4 of 5 transcripts intron retention (IR) between Exon 5 to 6 were reported (**Fig. 7a**). Presumably, the splice donor site is non canonical and can be skipped. **Fig. 7b** inspects the impact of Met267Ile substitution on properties of the protein, using conventional 3D protein metrics. The predicted protein characterization (with 12 properties such as protein stability, surface area, folding) were stable and uninterrupted relative to wild type ^24^). **Fig. 7c** shows multiple sequence alignment (MSA) of GLA candidate unannotated proteins. Note that the sequence of a protein variant (length 270 aa) resulted from skipping of exon-intron boundary, and resulted from extending reading of exon 5, with additional 3 aa, till reaching a stop codon. Collectively, these results highlight that the missense interpretation alone may not capture the variant’s pathogenic potential and disruption of transcripts due to mis-splicing is likely to explain the pathology of these ultrarare diseases. These findings demonstrate that MAGI offer a complementary perspective to classical protein-centric analyses. While traditional data support a missense effect on α-galactosidase A activity, MAGI uncover a concurrent splicing mechanism that likely contributes to disease pathology. This dual mechanism emphasizes the importance of integrating sequence-based splicing predictions with functional and clinical evidence, providing a more comprehensive understanding of how GLA c.801G>A drives Fabry disease.

**Figure 7.**
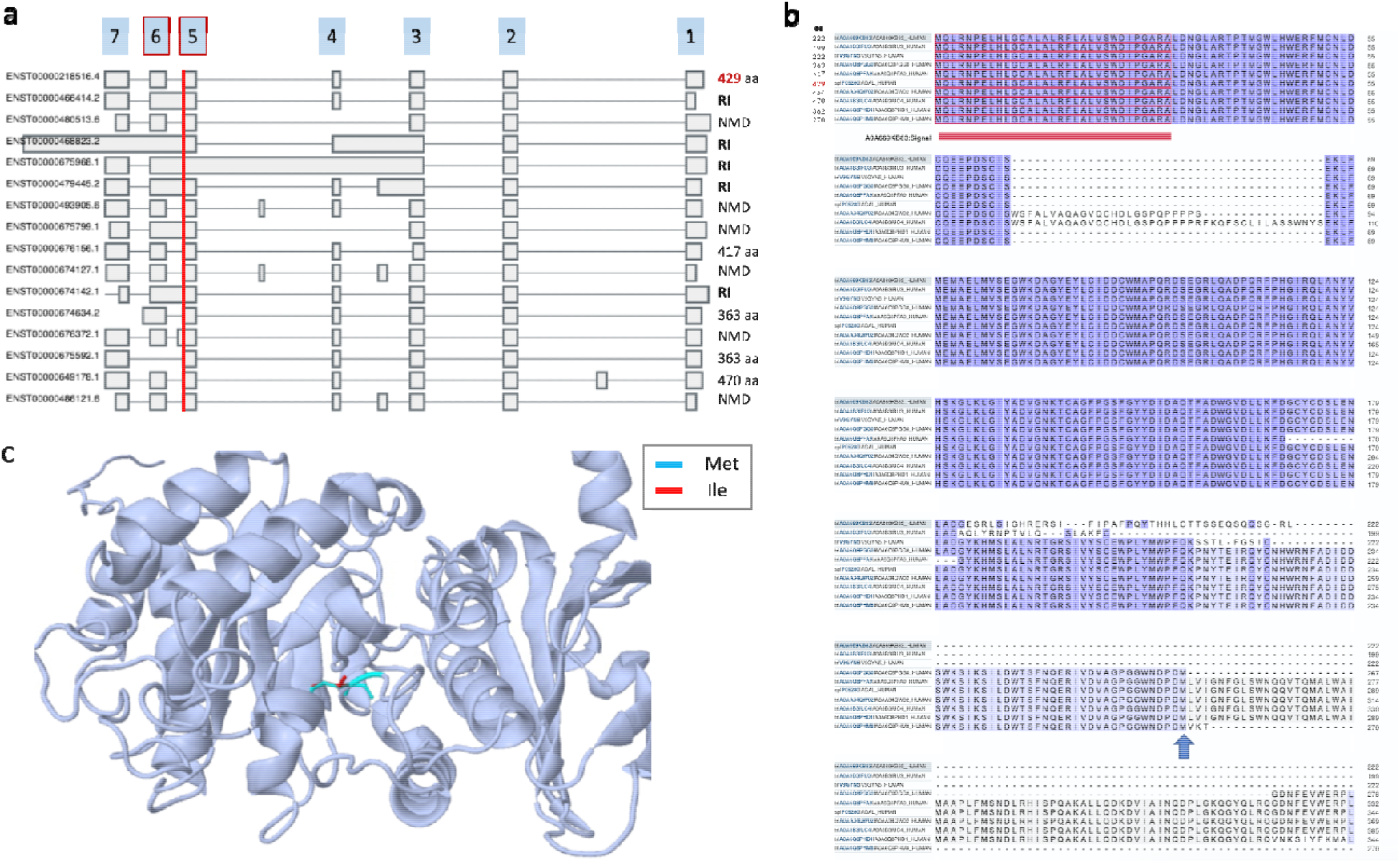
Multi-level characterization of the GLA p.Met267Ile variant. **(a)** Schematic representation of major transcript isoforms of the GLA gene (ENSG00000102393). Exons are shown as boxes and introns as connecting lines. Exon numbering is indicated above referring to the main protein variant based on RefSeq (NM_000169.3; UniProtKB-SwissProt P06280, 429 aa,). The red vertical line marks the position of the c.801G>A variant, located at the end of exon 5 and shared across multiple transcripts. **(b)** Multiple sequence alignment (MSA) of GLA proteins (α-galactosidase) across additional variants from the UniProt-TrEmbl database. The Met267Ile variant lies within a highly conserved domain at the end of an in-frame exon 5. At the top MSA, the protein length of each variant is indicated (in aa). The N-terminal signal peptide is highlighted (red box) and identical residues are shown in dark blue. **(c)** 3D structure of GLA protein (429 aa) from Missense 3D tool. Residue 267 is shown in stick representation, with the wild-type Met and the Ile substitution highlighted. The residue is located within the protein core.

### MAGI interpretation of VUS exposes an impact on chromatin state

The variant rs897804 (NM_013312.3:c.1464C>G; p.His488Gln; Chr19: 12766150-G>C) in the HOOK2 gene shows conflicting classifications in ClinVar, with one submission reporting it as pathogenic and two as benign. Overall, it is interpreted as having uncertain clinical significance (VUS).

The variant rs897804 (p.His488Gln), located in exon 15 of the HOOK2 gene, is predicted to have minimal impact on protein structure and function. Structural modeling based on PDB entry 4GKW indicates no disruption of any key structural features. Missense3D tool classifies this substitution as structurally neutral. Still, ClinVar assertions remain conflicting. At the population level, the variant is common (minor allele frequency = 0.32 in gnomAD). The SNP occurs in a poorly constrained genomic region (based on constrained coding region (CCR) percentile and low population-level tolerance score (pLI, see Methods). While these available scores argue against a deleterious protein-coding effect, the impact of rs897804 on function remains unsolved.

In contrast, MAGI-derived profiles highlight localized perturbations in regulatory features at the variant locus. The variant coincides with signals associated with open chromatin and regulatory activity (**Fig. 8a**), consistent with the presence of exonic cis-regulatory elements embedded within an accessible genomic region (**Fig. 8b**). The presence of regulatory elements in the vicinity of the SNP includes features linked to chromatin accessibility and transcript regulation, suggesting that rs897804 may alter regulatory grammar rather than protein structure. This interpretation provides a mechanistic basis for its association with complex traits, emphasizing that coding variants can exert functional effects through chromatin-mediated and transcript-level regulation rather than direct protein alteration.

**Figure 8.**
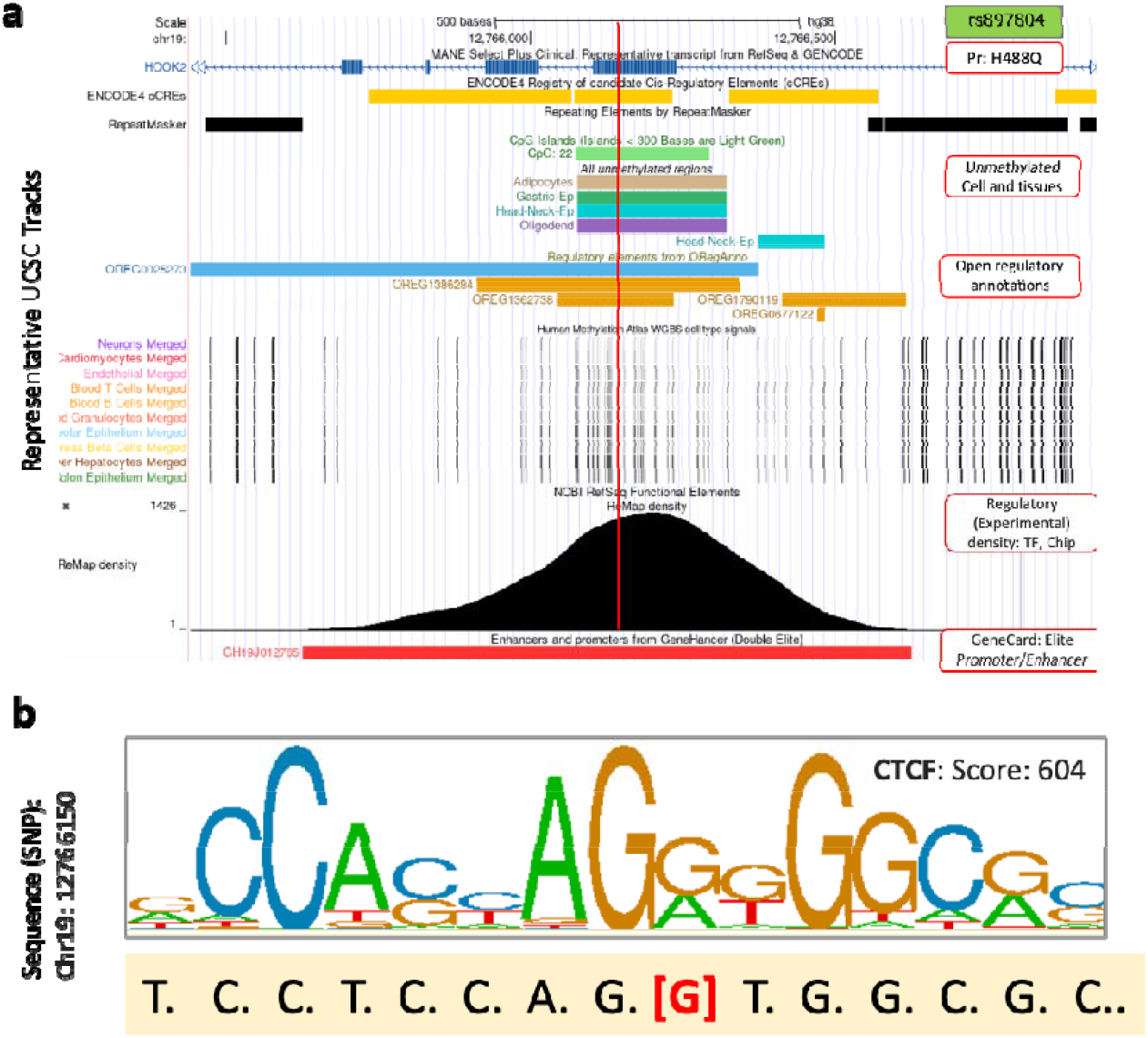
Track-based evidence for chromatin-associated regulation of rs897804. **(a)** The missense variant rs897804 (p.His488Gln), located in exon 15 of 22 of the HOOK2 gene. Several regulation tracks in a short region of 1500 nt centered around rs897804 suggest a dense regulatory region which is hypomethylated in specific tissues and cell types and includes enhancers and high density of cis transcription factor binding sites. **(b)** A short sequence around rs897804 matches the element that specifies CTCF binding sites. The WebLogo for CTCF element covers the SNP (G, red font). A CTCF element with statistical significance with p value of ~1e-05. The vertical red line indicates the position of the rs897804 SNP.

### MAGI generalization across species

We applied the framework to 580 curated Mendelian disease variants from the Online Mendelian Inheritance in Animals (OMIA) database ^22^, for SNPs and Indels across multiple non-human species. Examples were shown for dog (n = 360), cat (n = 185), and chicken (n = 35). All variants are curated with documented phenotypic effects. While epigenomic BigWig tracks are available only for humans, all 580 variants were successfully processed. MAGI performance for OMIA data is presented in Supplementary **Fig. S1**.

## Discussion

The central conceptual advance of MAGI is the transition from variant scoring to variant explanation. Most existing computational frameworks reduce variant interpretation to a short label of pathogenicity estimate, irrespective of whether the underlying perturbation involves coding disruption, altered transcript processing, chromatin remodeling, tissue-specific regulatory rewiring, failure in regulation etc ^25^. Recall that most of these algorithms and tools performed best to coding genes and their performance drops across regulatory regions ^26, Wang, 2023 #40, 27^. While such approaches have transformed large-scale variant prioritization, they remain fundamentally limited in their ability to provide mechanistic hypotheses that are clinically interpretable and experimentally actionable ^28^. MAGI addresses this gap by integrating genome foundation model (NTv3) functional track outputs ^28, 29^ with deterministic biological logic to produce explicit molecular consequences rather than abstract probability scores.

Recent years have seen rapid progress in genomic foundation models and deep variant effect predictors, including SpliceAI, Enformer, AlphaMissense, EVE, and NT-based architectures. These methods substantially improved predictive performance by leveraging evolutionary constraints, sequence context, or latent representations learned from large-scale genomic data. However, even the most accurate predictors generally compress diverse molecular effects into a single pathogenicity metric. MAGI differs conceptually from existing variant effect predictors in several respects. First, it is mechanism-centric rather than classification-centric, prioritizing biological explanation over pathogenicity scoring. Second, it integrates orthogonal genomic signals spanning exon structure, ORF integrity, splicing, chromatin state, tissue-specific transcription, and sequence-constrained regulatory motifs. Third, its deterministic inference layer produces explicit molecular rationales that can be inspected and experimentally tested. This approach aligns with recent efforts to improve foundation-model interpretability through sparse autoencoders (SAEs), which decompose latent representations into human-interpretable features ^30, 31^. Such transparency is particularly important in clinical genomics, where interpretability is increasingly required for diagnostic adoption and medical decision-making.

For validation, we challenge the full explanation associated with ClinVar and assess the mechanistic concordance with the magnitude of the model-derived perturbation signal. This observation suggests that genomic foundation models encode biologically meaningful representations extending beyond statistical sequence regularities ^19^. Variants with the strongest MAGI signals consistently aligned with curated expert interpretations, particularly for splice-site disruption, start-loss events, and ORF perturbations. Importantly, these categories are not merely associated with elevated pathogenicity scores, but correspond to well-defined molecular mechanisms ^32^. This distinction is fundamental because it suggests that the latent representations learned by genomic foundation models may already encode biologically interpretable structures that can be computationally extracted and linked to mechanistic variant effects.

Our results also highlight a persistent challenge in variant interpretation that concerns the incomplete penetrance and mechanistic heterogeneity. By integrating allele frequency with mechanistic burden, MAGI exposes variants that are molecularly disruptive yet tolerated at the population level. These “exonerated” variants likely represent cases involving compensatory regulation, tissue restriction, environmental dependence, or reduced penetrance. Conventional pathogenicity predictors often overestimate the clinical relevance of such variants. MAGI focus on molecular effect as a separated property relative to clinical-oriented disease consequence.

In this study we inspected several cases with the goal to illustrate MAGI interpretability and capacity to address complex impact of variants. The MAT1A and GLA case studies further illustrate the importance of transcript-aware interpretation. In both examples, conventional protein-centric interpretation was insufficient to explain the observed pathogenicity. MAGI identified concurrent transcript-level perturbations, including altered splice donor activity, intron retention, and isoform imbalance, thereby exposing dual molecular mechanisms. These findings are particularly relevant because many clinically interpreted missense variants are likely to exert composite effects that extend beyond amino acid substitution alone. Increasing evidence from multiplexed functional assays and transcriptomic profiling suggests that coding variants frequently perturb splicing ^33, 34^. While this phenomenon is well established for synonymous and canonical splice-site variants, it remains comparatively underappreciated for apparently disruptive missense mutations, where transcript-level effects may coexist with or even dominate protein-level consequences ^35, 36^. MAGI provide a framework capable of capturing these multi-layered effects (e.g., splicing, RNA stability, regulatory grammar) simultaneously and in a unified manner.

An additional strength of MAGI is its applicability beyond human clinical genomics. Despite being trained primarily on human-derived functional tracks, the framework generalized to curated Mendelian variants across multiple species. This observation suggests that mechanistic representations learned by genomic foundation models may capture sufficiently conserved principles of gene regulation and transcript organization to enable cross-species interpretation. Such portability may become increasingly important for comparative genomics, veterinary genetics, and functional annotation in organisms lacking extensive experimental resources.

Several limitations should nevertheless be acknowledged. First, MAGI remain dependent on the quality and completeness of the underlying genomic tracks and annotations. While we included 3602 tracks (BigWig) and 21 from BED files. We observed that most of the BigWig tracks are sparse and their contribution is quite limited. Refining the weights associated with the different tracks are expected to improve MAGI’s discovery rate. A known limitation concerns the shortage in validated regulatory regions and transcript models, particularly in non-coding regions and underrepresented tissues. Additionally, although deterministic logic improves interpretability, rule-based systems inevitably simplify biological complexity and may fail to capture nonlinear interactions between regulatory layers. Finally, the concordance for indels remained substantially lower than for SNVs, emphasizing the continued difficulty of modeling larger sequence perturbations using current foundation-model representations. Obviously, the use of ClinVar with >2 submitters and the associated rationales as a benchmark, introduces ascertainment biases toward known and well-studied mechanisms.

Increasing evidence from multiplexed functional assays and transcriptome-wide analyses demonstrates that many coding variants exert their effects through altered RNA processing and exon recognition rather than exclusively through changes in protein structure ^37, 38^. Although this mechanism is widely recognized for synonymous and canonical splice-site variants, transcript-level disruption remains comparatively underexplored for missense mutations that are typically interpreted primarily through protein-centric frameworks. Future work should integrate MAGI with experimental perturbation datasets, including MPRA, CRISPR saturation editing, long-read transcriptomics, and single-cell functional assays. Incorporating haplotype context across diverse populations and allele-specific regulation may further improve interpretation of complex and incompletely penetrant variants. More broadly, extending foundation-model inference to dynamic cellular states and chromatin organization could enable deeper mechanistic interpretation across regulatory scales. We anticipate that future clinical interpretation frameworks will increasingly combine probabilistic foundation models with explicit biological reasoning to move beyond predictive scoring toward mechanistic genomic inference.

## Methods

### Model and feature extraction

MAGI uses the Nucleotide Transformer v3 650M-parameter model (NTv3_650M_post). For each variant, 64-kb windows (GRCh38 coordinates) are processed to extract the reference and alt predictions across: (i) 21 structural genomic elements (BED tracks: coding, splice, regulatory, UTR, start/stop codon, ORF and other regulatory elements) as softmax probabilities, and (ii) 3,602 epigenomic tracks (BigWig; filtered from 7,362 to retain histone ChIP-seq, DNase-seq, CAGE, GTEx and RNA-seq tissue specific expression, full list in codabase). Per-track disruption is Δ_j_=P_j_(alt)−P_j_(ref).

### Sequence model features

The log-likelihood ratio LLR=log(P(Alt)/P(Ref)) quantifies sequence constraint and learned “predictability”, for SNPs. KL divergence between Ref and Alt token distributions captures local perturbation magnitude. Embedding-space distances (based on cosine, L2) are additionally computed and apply equally to indels.

### Impact scoring

Feature deltas are aggregated: MAGI(v)=∑_j_ log(1+⍰z_j_ (v) ⍰), where z _j_ (v) = ( Δ _j_ (v) − μ_j_) / σ_j_. Background parameters (μ_j_,σ_j_) estimated from ClinVar benign-classified variants.

### Mechanistic interpretation

Disruption signals are mapped to mechanistic annotations via deterministic, hierarchical rules prioritizing structural elements, then epigenomic tracks, then sequence-model metrics.

### Concordance evaluation

Concordance evaluation. NT-v3 mechanistic signals were compared against expert-curated ClinVar submission rationales for pathogenic and benign variants, with non-placeholder free-text explanations (9,786 SNPs and 12,301 indels after ≥2-star review filtering and de-duplication). For each variant, an LLM judge (Gemini 3 Flash Preview; temperature 0.1) received the aggregated ClinVar rationale together with summarized NT delta signals: top BED annotation changes, epigenomic/transcriptional track changes, and MLM sequence-context scores (plus indel embedding and KL metrics for indels). The judge assigned concordance as concordant (NT signals support the mechanism described in the rationale), partially concordant (partial support), discordant (signals contradict or fail to support the described mechanism), or not applicable (rationale describes purely structural protein effects with neutral NT signals). Variant-type-specific system prompts defined these categories and signal-scale interpretation rules (full prompts in codebase). A blinded manual review of 100 variants was performed to validate judge assignment.

## Data and code availability

The data used in this study are publicly available. Human SNP annotations and associated clinical rationales were obtained from the SongLab-ClinVar benchmark, while indel annotations were obtained from the ClinVar-GFM-Bench dataset, both of which are derived from the public ClinVar resource. Original ClinVar records, annotations and rationale are available through ClinVar (https://www.ncbi.nlm.nih.gov/clinvar/).

Animal variant data were obtained from OMIA (https://omia.org/). Population allele frequencies were obtained from gnomAD v4.1.1 (https://gnomad.broadinstitute.org/). Gene annotations are based on MANE Select v1.3. The NTv3 foundation model is available from Hugging Face (https://huggingface.co/InstaDeepAI/NTv3_650M_post). Source code for the MAGI pipeline, including inference, analysis, and interpretation modules, is available on GitHub at https://github.com/ddofer/magi. An interactive web application for multi species variant analysis is freely available at https://huggingface.co/spaces/GrimSqueaker/MAGI.

## Supplementary Figures

**Figure S1.**
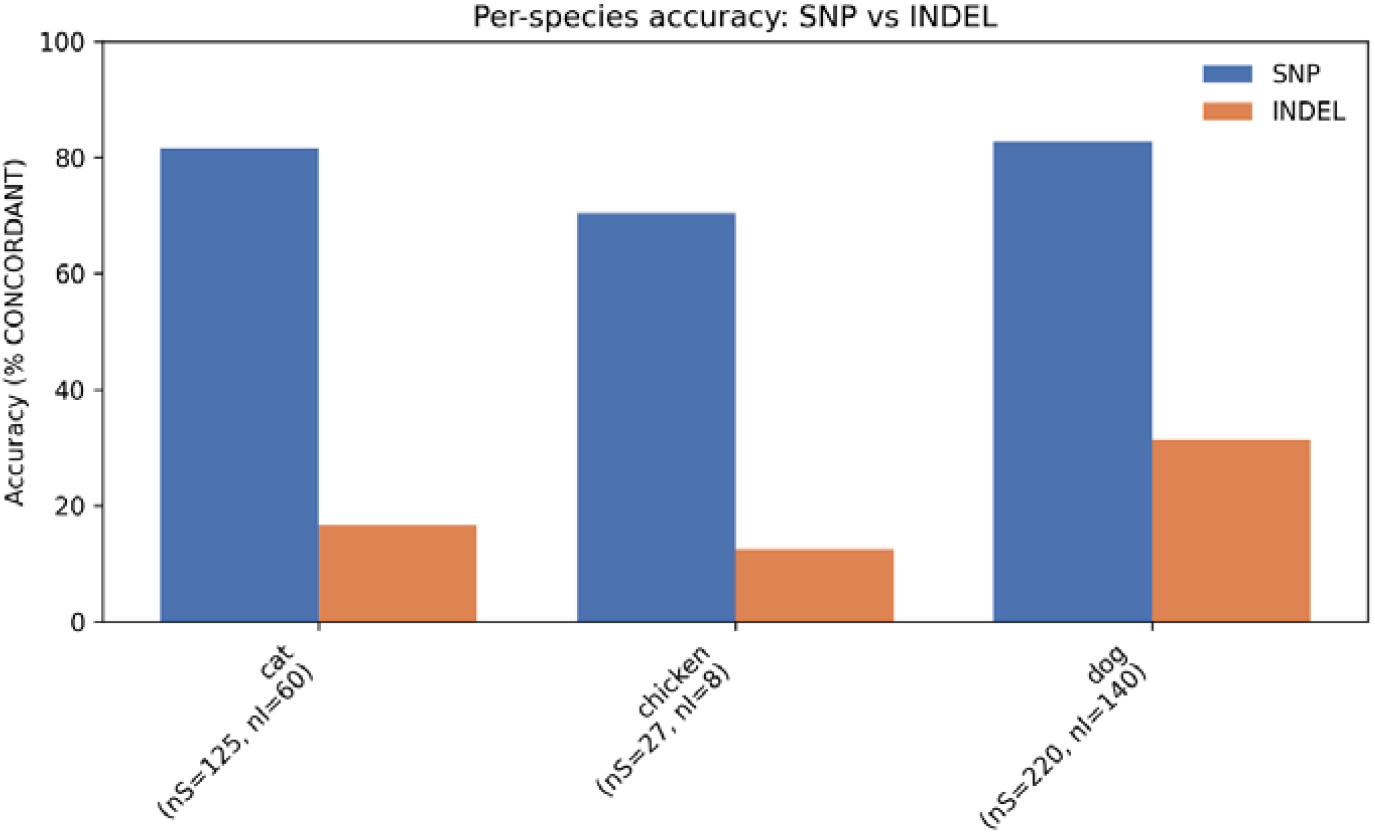
Semantic concordance with clinical with expert clinical rationales. Concordance is stratified by variant class (SNPs vs. Indels) and the listed mammals. Note the relatively poor concordance (%) associated with indels across species.

